# Interacting cortical gradients of neural timescales and functional connectivity and their relationship to perceptual behavior

**DOI:** 10.1101/2022.05.05.490070

**Authors:** Matthew J. Boring, R. Mark Richardson, Avniel Singh Ghuman

**Affiliations:** Center for Neuroscience at the University of Pittsburgh, University of Pittsburgh, Pittsburgh, PA, USA; Center for the Neural Basis of Cognition, University of Pittsburgh and Carnegie Mellon University, Pittsburgh, PA, USA; Department of Neurological Surgery, University of Pittsburgh, Pittsburgh, PA, USA; Department of Neurosurgery, Massachusetts General Hospital, Boston, MA, USA; Harvard Medical School, Boston, MA, USA

**Keywords:** Cortical gradients, functional connectivity, neurodynamics, neural timescales, intracranial electroencephalography

## Abstract

Cognitive acts take place over a large range of temporal scales. Numerous corresponding gradients in neurodynamic timescales and long-range cortical interactions are believed to provide organizational constraints to the brain and influence neural populations’ roles in cognition. However, it is unclear if gradients in various types of neural timescales and functional connectivity arise from related or distinct neurophysiological processes and if they influence behavior. Here, intracranial recordings from 4,090 electrode contacts in 35 individuals were used to systematically map gradients of multiple aspects of neurodynamics, neural timescales, and functional connectivity, and assess their interactions along category-selective ventral temporal cortex. Opposing functional connectivity gradients, with decreasing connectivity to visually responsive regions and increasing connectivity to regions that were not visually responsive, were observed along the ventral visual hierarchy. Endogenous neural timescales were correlated with functional connectivity to visually responsive regions after removing the effects of shared anatomical gradients, suggesting that these properties influence one another. Different stimulus evoked and endogenous timescales exhibited gradients with longer dynamics along the ventral visual hierarchy, but none of these timescales were significantly correlated with one another. This suggests that local neural timescales depend on neural and cognitive context and different timescales may arise through distinct neurophysiological processes. Furthermore, activity from neural populations with faster endogenous timescales and stronger functional connectivity to visually responsive regions was more predictive of perceptual behavior during a visual repeat detection task. These results reveal interrelationships and key distinctions among neural timescale and functional connectivity gradients that together can influence behavior.

## Introduction

A neural population’s functional properties, including its dynamics and its functional connectivity to other brain regions, are ultimately linked to that population’s role in cognition and perception. Several gradients in functional properties have been shown to exist along the cortical axis spanning from primary sensory/motor areas to association cortices (1, 2, 6–9). These gradients are thought to be related to key organizing principles of the cortex, guiding how different regions contribute to cognition and perception (2, 7, 10, 11). For example, there are gradients in the timescales over which neural populations process information and endogenously fluctuate along this axis, with longer timescales further along cortical hierarchies (5, 8–10, 12–15), though it remains unclear if stimulus processing and endogenous timescales are related to one another. Gradients of functional connectivity are also seen along this axis, with decreasing unimodal connectivity and increasing transmodal connectivity along cortical hierarchies (2, 16). These network-level neural properties likely influence local timescales, other computational characteristics of neural populations, and these populations’ relationship to behavior (2, 6, 7, 17). However, empirical evidence linking functional gradients in local dynamics with gradients in the long-range connectivity of neural populations is limited. Additionally, it is uncertain to what degree these gradients relate to a neural population’s role in behavior.

One prevalent functional gradient in cortex is the increasing timescales over which neural populations integrate information when moving from primary sensory/motor to association cortices (5, 8, 9, 12, 13, 18–20). For example, rapidly varying acoustic inputs represented in low-level auditory cortex are combined into more complex representations in higher order auditory cortex, which operates over longer timescales (14). These neural timescales, or temporal receptive windows, are related to the rate of decay of representations within neural populations (5, 12, 13, 18), because longer decay rates allow for more pieces of information to be integrated into a single representation. Stimulus-unrelated, endogenous timescales also lengthen along this axis, measured through the temporal autocorrelation of neural activity (5, 7, 8, 10, 21). It is unclear the extent to which stimulus-related and endogenous timescales relate to one another.

Another key aspect of neural dynamics, which is less well understood, is information processing dynamics, including the initial rate at which neural populations discriminate between stimuli (i.e., the rise time of discriminant information in neural activity). These information processing dynamics relate to the speed of cortical computation and thus, ultimately limit the speed of decision and action processes (3). Despite the importance of a neural population’s information processing dynamics in cognition and perception, the functional characteristics that are associated with neural populations that processes information more quickly or slowly remain unclear (12, 13, 19).

In addition to anatomical gradients in local neural dynamics, opposing anatomical gradients in long-distance connectivity to association versus primary sensory/motor cortices have also been demonstrated in human cortex. Unimodal connectivity, primarily within sensorimotor regions, decreases when moving up cortical processing hierarchies while transmodal connectivity linking multiple sensory domains increases (1, 2). However, it is unclear how gradients in local dynamics interact with gradients in long-range functional connectivity. In silica, circuit models of cortical processing suggest that inter- and intra-areal connectivity patterns help constrain a neural population’s timescale (6, 11, 22), which has received some support from low temporal resolution measures of brain activity (17, 23).

Finally, the functional properties that constrain a neural population’s dynamics and long-range cortical connectivity ultimately constrain how that population contributes to cognition and perception. However, open questions remain regarding which aspects of a neural population’s anatomical position, neurodynamics, or functional connectivity is related to its ability to predict behavior.

In the current study, category-selective neural populations in ventral temporal cortex (VTC) were used as a model to examine the relationship between anatomical gradients in local cortical processing and long-range cortical interactions. We also explored how information processing dynamics, endogenous timescales (i.e., neural dynamics not directly linked to the exogenous, stimulus-evoked response; which we operationalize using the prestimulus period when no stimulus was being presented), and long-range cortical connectivity interact with each other beyond any shared anatomical gradients, and which of these gradients were associated with the ability of a population’s activity to predict response time during single trials of a visual 1-back task.

## Results

Activity was recorded from 1,955 VTC electrode contacts (out of a total of 4,090 intracranial electrode contacts distributed throughout the brain) in 35 patients with pharmacologically intractable epilepsy (*Fig. S1*) as they viewed images of objects (face, body, word, hammer, house, or phase scrambled image) during a 1-back task. Multivariate Naïve Bayes classifiers were used to predict the category of object participants were viewing during individual trials of the task using sliding 100 ms windows of single trial potentials (stP) and single trial high frequency broadband activity (stHFBB) recorded from individual electrode contacts. At this stage of the analysis, these signal components were combined since previous studies have suggested that they contain complementary information (24), though in further analyses they were examined separately. Out of the 1,955 VTC electrode contacts, activity recorded from 390 electrode contacts (mean = 11; SD = 14 electrode contacts per patient) could reliably predict (p < 0.001, corrected via permutation testing) which category participants were viewing during single trials of the task (*Fig. 1*). Information processing dynamics were estimated from the activity of the neural populations recorded by these contacts. Specifically, the time-course of category-discriminant information processing in these category-discriminant neural populations was calculated by computing the mutual information (in bits) between the classifier outputs and the true category labels. The functional properties of these populations were computed to examine the relationship between these variables and anatomical axes of VTC (see *Methods*). Specifically, we examined gradients of, and interactions between, nine factors: two stimulus response timescales (factors 1 and 2): initial rise duration and maintanence of category-discriminant information (see *Fig 2A* for illustration); category-discriminant information onset time and peak magnitude (factors 3 and 4; see *Fig 2A* for illustration); two endogenous (prestimulus) timescales (factors 5 and 6): the timescale of decay, “tau”, for the prestimulus stP and stHFBB autocorrelation functions (see *Fig 4A* for illustration); functional connectivity to visually responsive populations and to populations that were not significantly visually responsive (factors 7 and 8); and the accuracy of a neural population’s activity for predicting behavioral response time (RT; factor 9).

**Fig. 1.**
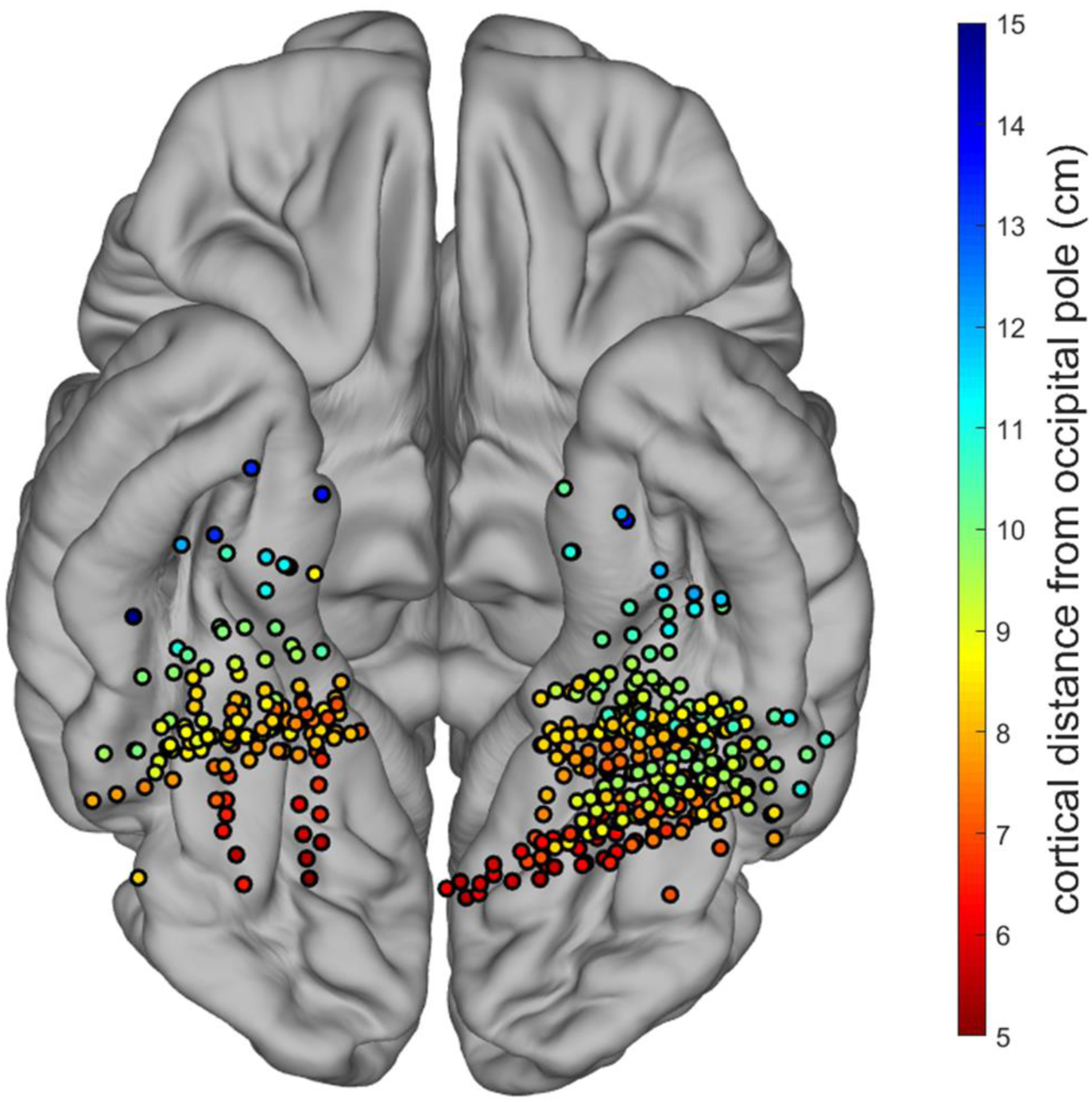
Spatial topography of electrode contacts recording from neural populations that achieved peak category-discriminant information greater than chance at the p < 0.001 level corrected for multiple temporal comparisons. Colors represent the electrode contact’s 5 cortical distance from the occipital pole calculated using subject-specific anatomy. Depth electrode contacts are plotted below the cortical surface for clarity. Contacts that appeared to be outside of the MNI standard brain due to differences in individual brain sizes were projected to the nearest MNI cortical vertex in this figure solely for illustrative purposes. The proportion of left vs. right hemisphere category-discriminant contacts was comparable to the proportion of total left vs. right hemisphere VTC implants (see *Supplementary Text*).

**Fig. 2.**
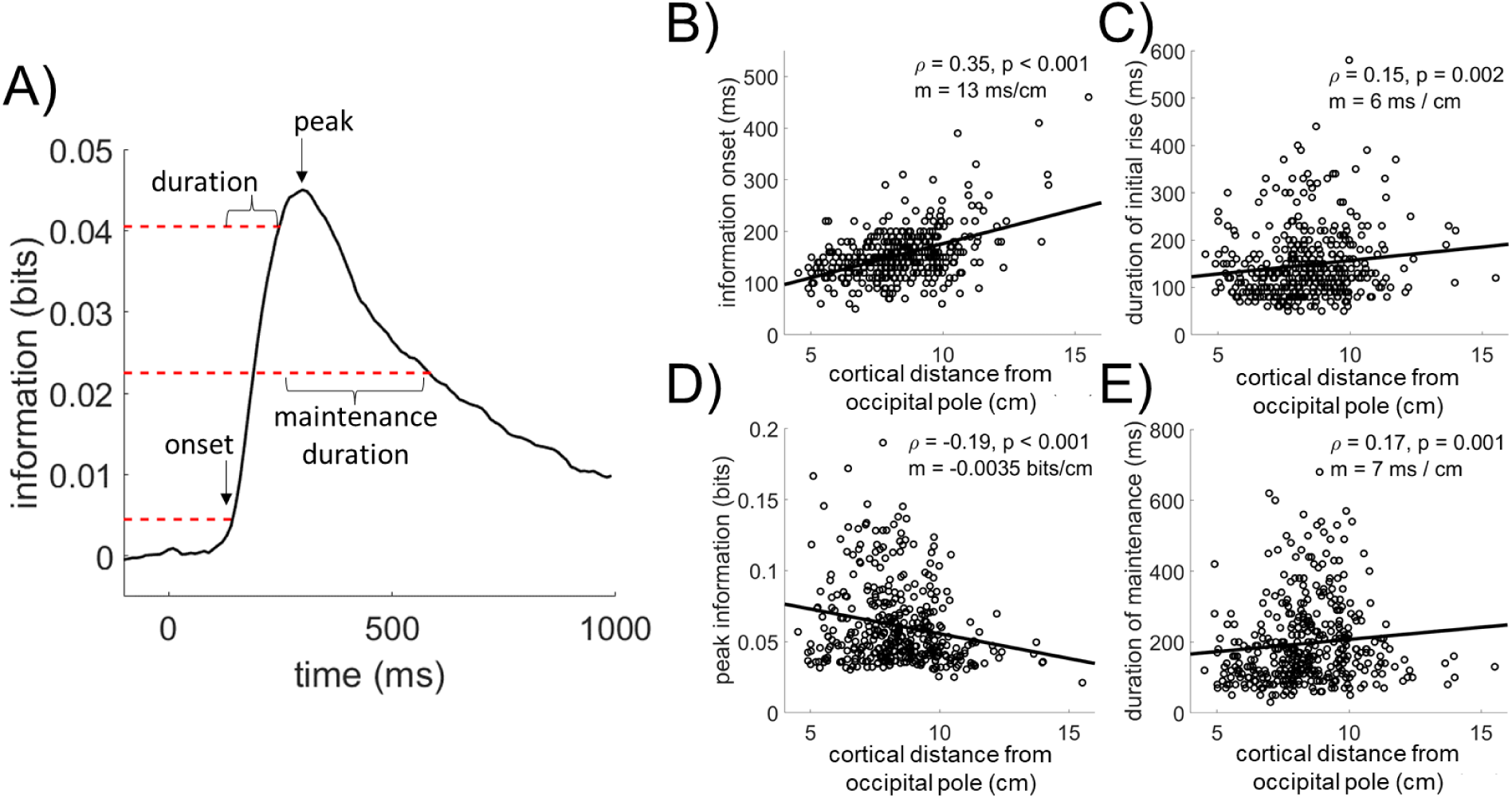
Category-discriminant information processing gradients along the ventral visual hierarchy. A) The time-course of category-discriminant information processing was computed for each neural population. The average time-course across category-discriminant VTC populations is illustrated here. From each neural population’s information processing time-course, the information onset time (panel B), processing duration (C), peak magnitude (D), and maintenance duration (E) were computed. Simulations confirmed that decreases in information amplitude and information processing duration are independent using our methods (*Fig. S6*). B) The onset of category-discriminant information, defined as the timepoint the information reached 10% of the maximum before peaking, was significantly correlated with the position of that neural population along the ventral visual hierarchy. The black line indicates the least-squares regression fit. Spearman’s ρ and associated p-value shown on top right (n = 390). Spearman correlation was used because it is both more robust to outliers relative to Pearson’s and is sensitive to non-linear monotonic relationships between variables, though this also means that the line drawn is not representative of the ρ. Slope of the least-squares regression line (m) indicated a 13 ms per centimeter increase in onset latency moving along VTC. Information onset was significantly associated with distance along the visual hierarchy even after correcting for cross-patient differences in onset latency (T(388) = 7.20, p < 0.001, tied-rank mixed-effects model). C) The duration of the initial rise in category-discriminant information, defined as the time between the onset of information and the time it took the population to reach 90% of its peak information, was negatively correlated with distance along the visual hierarchy. The 90% threshold is used for the peak time because it better captures the initial rise in cases where there is a shallow peak among an extended plateau in the discriminant information time-course. *<u>Note</u>*: All correlations remain significant across a substantial range of the heuristic thresholds chosen to define them (*Fig. S7*), thus the selection of 10% and 90% as thresholds for onset and peak time do not drive these effects. The slope of the least-squares regression line indicated a 6 ms increase in the duration of the initial rise of information per cm of VTC. This relationship did not reach p < 0.05 when correcting for random cross-patient effects (T(388) = -1.55, p = 0.12, tied-rank mixed-effects model). D) Peak category-discriminant information was negatively correlated with distance along the visual hierarchy, with a decrease of -0.0035 bits/cm. This relationship did not reach p < 0.05 when correcting for random cross-patient effects (T(388) = -1.62, p = 0.11, tied-rank mixed-effects model). E) Information maintenance duration, defined as the time between when the information first reached 90% and the time when it first decayed to 50% of the peak, was positively correlated with distance along the visual hierarchy. The slope of the least-squares regression line indicated a 7 ms increase in the duration of maintenance of information per cm of VTC. This relationship trended to p < 0.05 significance when correcting for random cross-patient effects (T(388) = 1.87, p = 0.063, tied-rank mixed-effects model).

### Gradients of Information Processing Dynamics

The cortical distance from the occipital pole, which roughly corresponds to the fovea in primary visual cortex, was used to approximate the position of neural populations along the hierarchical axis of the ventral visual stream (25). Distance along this axis was correlated with several aspects of information processing in these category-discriminant neural populations (*Fig. 2; see Fig S2* for an example from a single subject). Along this axis, neural populations demonstrated increasing onset latencies and increasing durations of their initial rises in category-discriminant information. Additionally, neural populations maintained category-discriminant information longer after peaking, despite reaching smaller peak magnitudes, when moving along the visual hierarchy. See *Fig S6* for simulations demonstrating the independence of peak magnitude and rise duration metrics.

In addition to examining discriminant information, we also examined the dynamics of the non-discriminant neural responses. Specifically, gradients in category-indiscriminant visual responses (discriminating all categories from baseline rather than categories from one another as in *Fig. 2*) in the same neural populations were examined (*Fig. 3*). Populations demonstrated increasing onset latencies and decreasing peak magnitudes of visual responsiveness when moving along the ventral visual hierarchy, like the gradients observed for category-discriminant information. However, there was no comparable increase in the duration of the initial rise in visual responsiveness along this axis and visual responsiveness was maintained for shorter durations in populations further along the visual hierarchy, which was opposite of the gradient observed for category-discriminant information. The contrast between visual response dynamics and category-discriminant information processing dynamics highlight differences in the neural encoding of these two levels of stimulus information (26, 27).

**Fig. 3.**
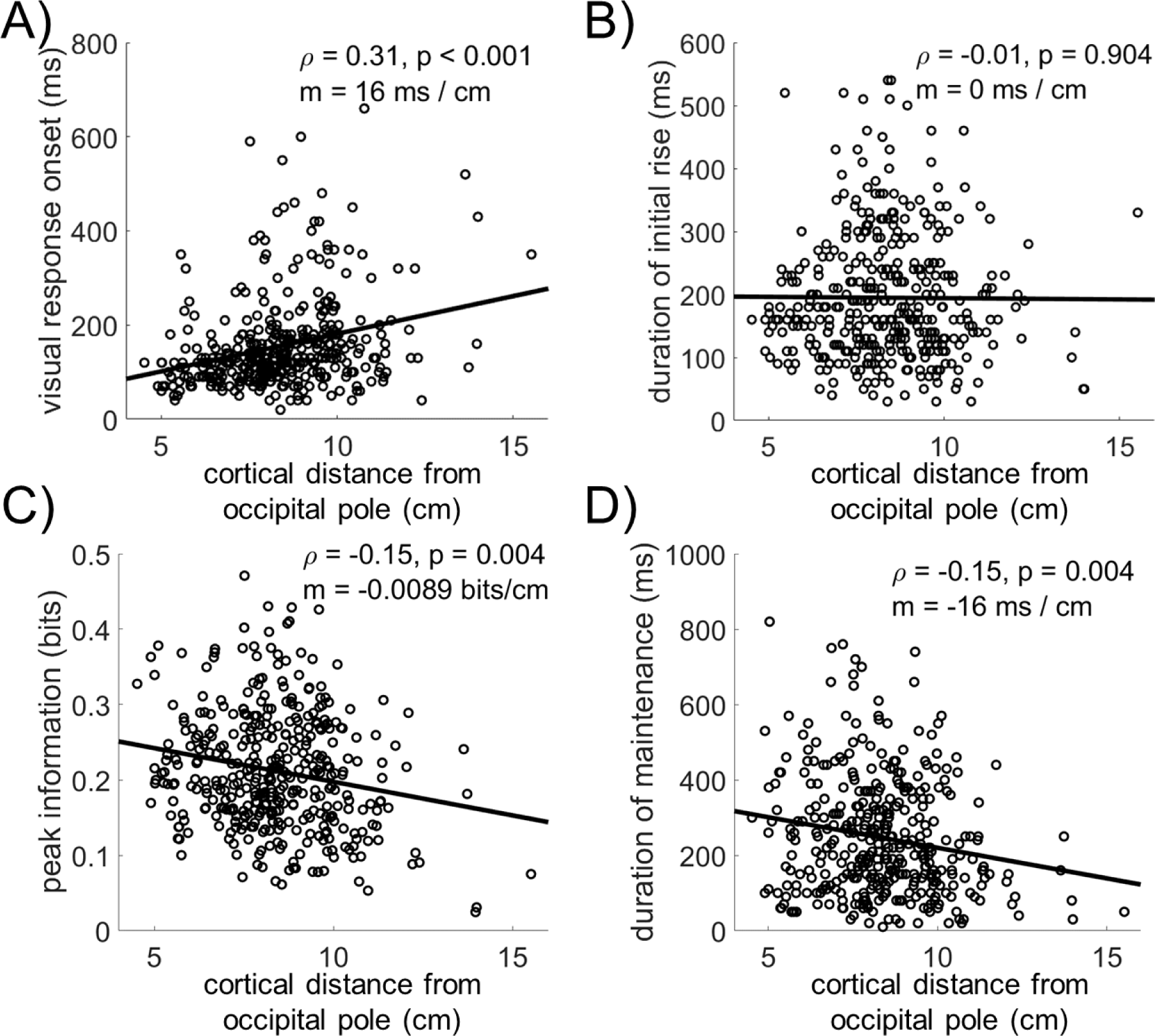
Visual response dynamics along the ventral visual hierarchy. Visual response dynamics were extracted by classifying all stimulus categories versus baseline with similar classifiers used to extract category-discriminant information (*Fig. 2*), making this a measure of the effect size between visual response versus baseline. A) Onset of the visual response was positively correlated with a neural population’s distance along the visual hierarchy. This effect held when correcting for random cross-patient effects (T(388) = 6.23, p < 0.001, tied-rank mixed-effects model). Onset latency of the visual response was not significantly different than the onset of category-discriminant information (T(389) = 0.11, p = 0.91, paired T-test). B) Duration of the initial increase in visual responsiveness was not significantly correlated with distance along the visual hierarchy, unlike the significant positive correlation observed for category-discriminant information (*Fig. 2C*). C) Peak magnitude of visual responsiveness was negatively correlated with distance along the visual hierarchy. This effect held when correcting for random cross-patient effects (T(388) = - 2.26, p < 0.001, tied-rank mixed-effects model). D) Visual response maintenance duration was also negatively correlated with distance along the visual hierarchy, which held when correcting for random cross-patient effects (T(388) = 5.45, p < 0.001, tied-rank mixed-effects model). This was opposite of the relationship between information maintenance duration and distance along the visual hierarchy observed for category-discriminant information (*Fig. 2E*).

### Gradients of Endogenous Neural Timescales

Next, the endogenous neural timescales of VTC populations were quantified by computing the autocorrelation of prestimulus activity at multiple temporal lags and modelling the resulting autocorrelation function with an exponential decay function (5, 22; *Fig. 4*). When moving along the visual hierarchy, neural populations demonstrated increasing time-constants of decay (tau) in the autocorrelation function of their prestimulus stP, indicating that their activity exhibited longer timescales/slower dynamics along this axis. This is consistent with previous studies observing slower endogenous timescales when moving up sensory processing hierarchies (2, 5, 7, 18, 19, 28). Conversely, neural populations demonstrated shorter timescales in their prestimulus stHFBB activity when moving along the ventral visual hierarchy. Time-constants across stP and stHFBB signal components were not significantly correlated with one another across electrode contacts (ρ(390) = -0.05, p = 0.33), highlighting the differentiability of these two aspects of the neural signal (29–31). These results show that these components of the endogenous neural activity demonstrate distinct timescales that have opposite gradients along the ventral visual hierarchy.

**Fig. 4.**
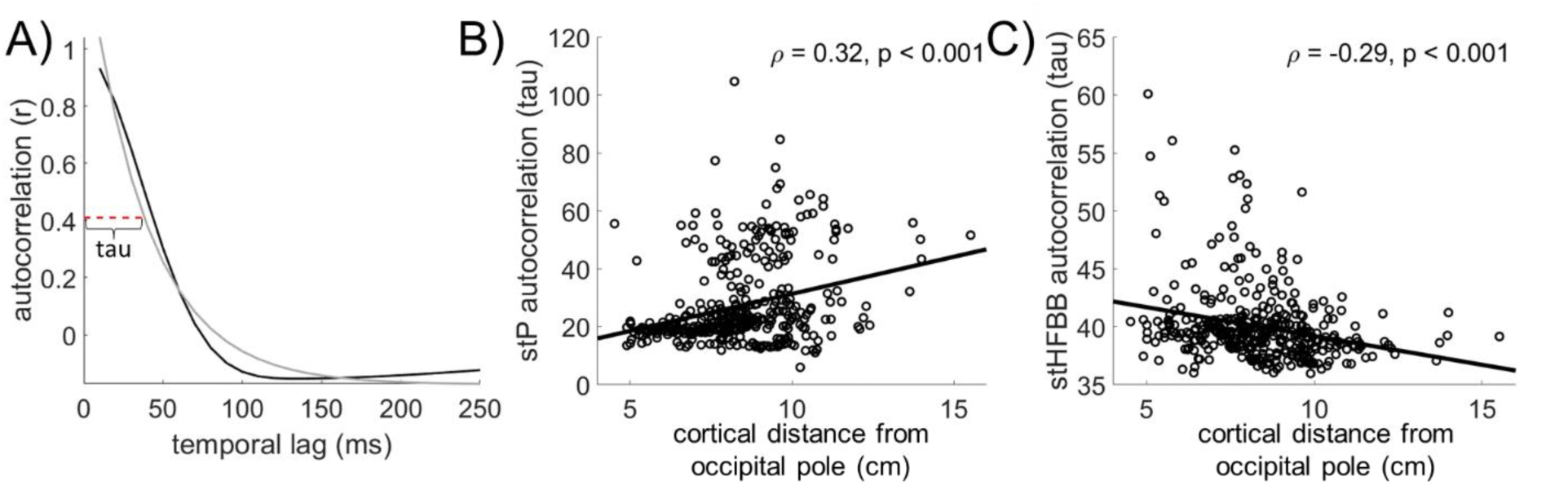
Prestimulus neural timescales along the ventral visual hierarchy. A) For each neural population, the autocorrelation function during the -500 to 0 ms prestimulus period was computed for temporal lags ranging from 1 to 250 ms, averaged across trials (black line is the average across populations), and fit with a single exponential decay function (gray line). The timescale (tau) indicates how fast the fitted exponential function decays (red dashed line; computed like those in (5)) and was correlated with other functional properties of the category-discriminant neural populations’ activity. B) The autocorrelation function of single trial potentials (stP) decayed more slowly when moving up the visual hierarchy, indicating that stP in more anterior VTC had higher autocorrelations at greater lags (longer timescales) relative to more posterior neural populations. This relationship held when correcting for random cross-patient effects (T(388) = 8.03, p < 0.001, tied-rank mixed-effects model). C) The autocorrelation function of single trial high frequency broadband (stHFBB) decayed more quickly when moving up the visual hierarchy, indicating that stHFBB in more anterior VTC had lower autocorrelations at greater lags (shorter timescales) relative to more posterior neural populations. This relationship also held when correcting for random cross-patient effects (T(388) = -5.32, p < 0.001, tied-rank mixed-effects model).

Gradients in both information processing dynamics and neural timescales were present in individual patients (*Fig. S2*) and several generalized across patients (linear mixed-effects models *Fig. 2 & 3* captions). The gradient of information processing onset was stronger in the left compared to the right hemisphere (*Supplementary Text*). Notably, neural populations selective for individual categories demonstrated different gradients in neural dynamics relative to those selective for other categories along the ventral visual hierarchy, with face selective populations generally displaying shallower posterior-anterior gradients (*Supplementary Text, Fig. S3, Table S1*). Given the differences in prestimulus neural timescales exhibited in stP and stHFBB, we recomputed gradients in information processing dynamics from these signal components separately. With a few notable exceptions, stimulus related information processing dynamics demonstrated similar gradients for stP and stHFBB across these components when moving along the visual hierarchy (*Supplementary Text* and *Fig. S4*).

### Gradients of Functional Connectivity

After examining gradients in information processing and endogenous timescales, we examined gradients in functional connectivity along the ventral visual hierarchy. Specifically, a measure of functional connectedness to the rest of the brain, the average prestimulus phase-locking value (PLV), was calculated between the 390 category-discriminant VTC electrode contacts and all other electrode contacts implanted within the same patient (on average 115 electrode contacts, SD = 41; note that none of the results reported below change substantially whether functional connectivity was calculated during the prestimulus or the poststimulus period as the Spearman correlation between the prestimulus and poststimulus connectivity metrics was > 0.95). These “other” electrode contacts were located across the entire brain, not only in VTC (*Fig. S1*). Previous fMRI studies suggest opposite gradients in functional connectivity to unimodal sensory vs. association and transmodal areas when moving along sensory processing streams (1, 2). Therefore, we separately computed the functional connectivity of VTC category-selective contacts to visually responsive contacts (p<0.001, for visual response vs. baseline, corrected for multiple temporal comparisons) and to those that were not visually responsive. Also, given the wide variability of electrode coverage across patients, pooling connectivity across visually responsive and not visually responsive contacts allowed us to partially overcome this cross-patient anatomical heterogeneity.

Connectivity between VTC electrode contacts and visually responsive contacts decreased when moving up the visual hierarchy. In contrast, the connectivity between VTC contacts and contacts that were not significantly visually responsive increased when moving up the visual hierarchy (*Fig. 5*). Decreasing functional connectivity to visually responsive regions and increasing functional connectivity to regions that do not demonstrate strong visual responses is generally consistent with previous fMRI studies showing opposing anatomical gradients along VTC for functional connectivity to unimodal versus transmodal regions (2).

**Fig. 5.**
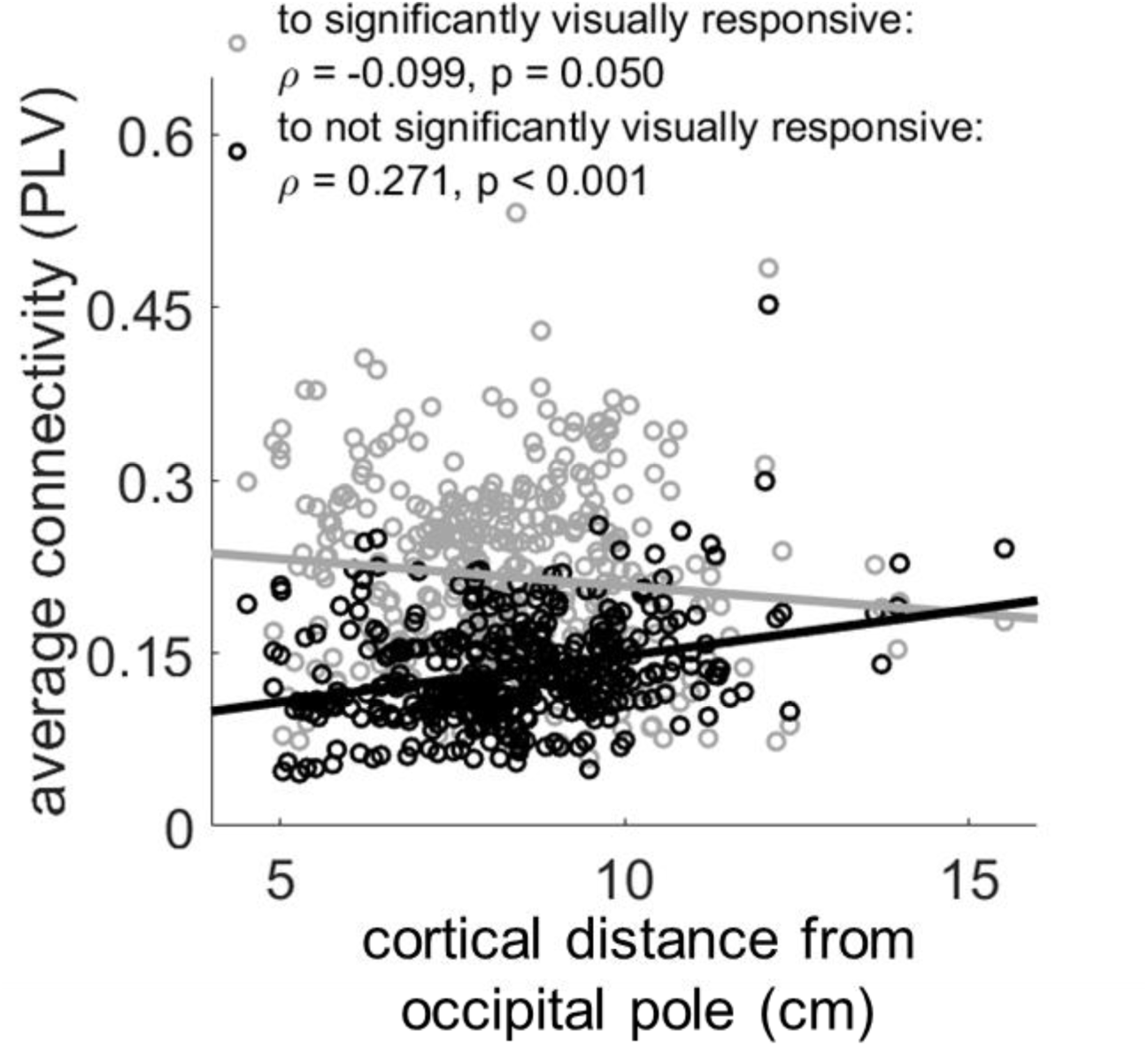
Gradients in long-range cortical interactions along the ventral visual hierarchy. The change in connectivity to visually responsive regions moving along VTC was opposite of the change in connectivity to populations that were not visually responsive. Connectivity to significantly visually responsive regions decreased along this axis, even when accounting for random cross-patient effects (T(388) = -4.42, p < 0.001, tied-rank mixed-effects model). On the other hand, connectivity to regions that were not significantly visually responsive increased along this axis, even when accounting for random cross-patient effects (T(388) = 3.98, p < 0.001, tied-rank mixed-effects model).

### Gradients of Ability to Predict Behavior

Additionally, we examined whether a neural population’s role in visual perceptual behavior exhibited an anatomical gradient. This was done by predicting the RT of patients, using sliding windows of neural activity recorded at each category-selective VTC electrode contact, during trials of the 1-back task where patients correctly responded that an object was presented twice in a row. How predictive the activity in an electrode contact was of RT was used as a measure of how much the activity from that neural population contributed to perceptual behavior. When considering stP and stHFBB together, the ability of a VTC neural population’s activity to predict RT was not significantly correlated with distance along the visual hierarchy (ρ(390) = 0.02, p = 0.75). However, when considering them separately, a neural population’s ability to predict RT decreased along the visual hierarchy when looking at stHFBB but increased when looking at stP. These differences highlight nuances in large-scale neuroanatomical gradients when considering different aspects of the neural signal (29–31).

### Relationships between and among neurodynamics, functional connectivity, and behavior

Given corresponding anatomical gradients in local dynamics and long-range cortical interactions, a key question is, to what degree these gradients are interrelated beyond shared anatomical axes. To explore this question, the partial correlations between these functional properties of category-selective VTC populations were calculated after removing the effects of distance along the visual hierarchy (*Fig. 6*). Note that Spearman’s partial correlation was used to remove any monotonic relationship to distance along the visual hierarchy, not only linear relationships (see *Fig. S5* for the non-partialed correlations).

**Fig. 6.**
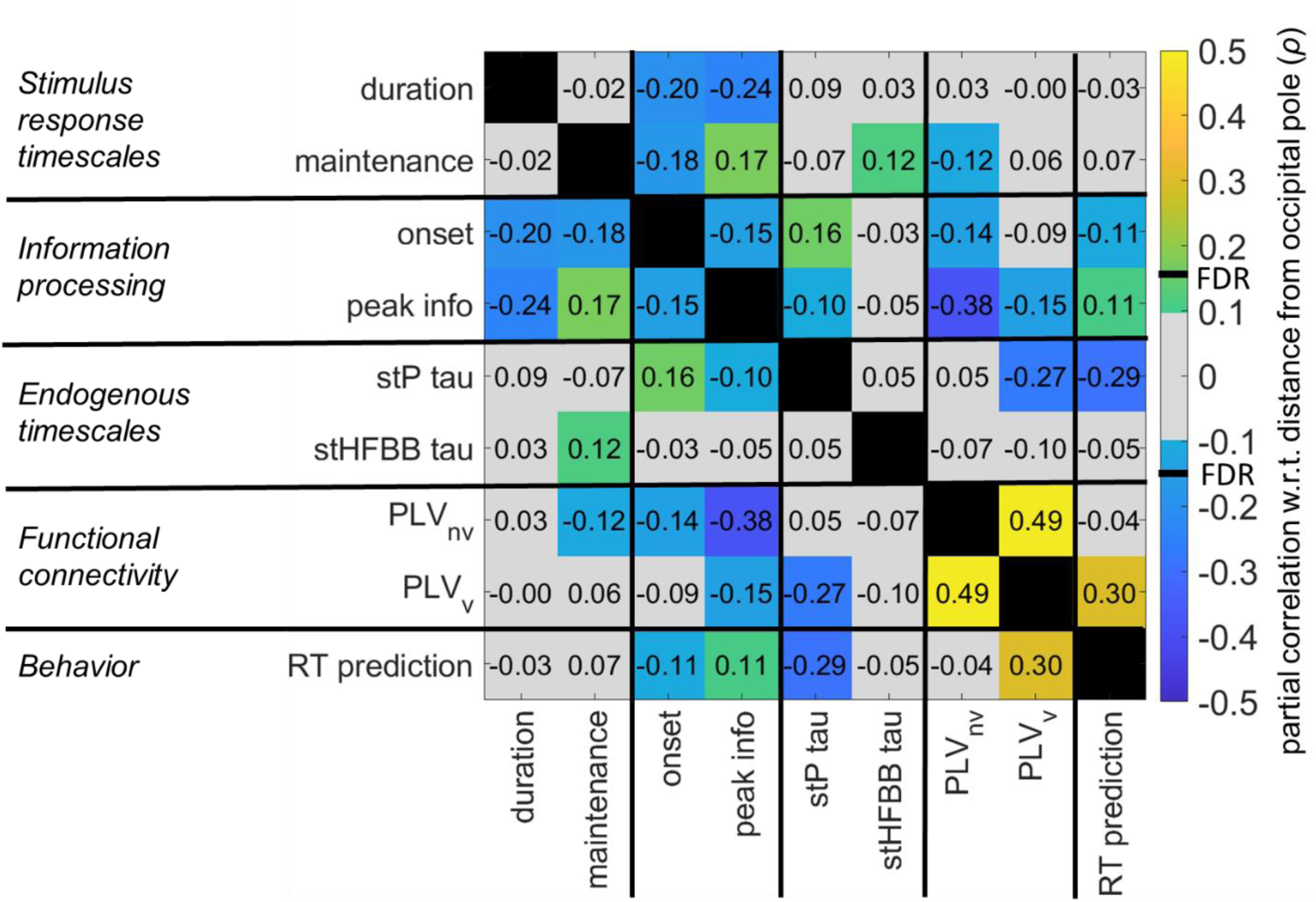
Interactions between local dynamics, long-range cortical interactions, and behavioral correlations. Partial correlation matrix between local dynamic properties and long-range cortical interactions after removing the effect of cortical distance along the visual hierarchy (see *Fig. S5* for non-partialed correlations). Colored squares are significant at the p < 0.05 level (uncorrected). The false-discovery rate adjusted critical value corresponds to ρ = ±0.146. Within each square is the partial Spearman correlation coefficient for the variables in the corresponding row and column. The matrix is symmetric across the diagonal. Several properties of the local information processing dynamics, including information onset, peak magnitude, duration of the initial rise, and the amount of time the information was maintained, were significantly correlated to one another besides sharing a common anatomical gradient. The partial correlation between peak information and functional connectivity was also significant after removing the effect of distance along the visual hierarchy. The partial correlation between neural timescale (stP tau) and connectivity to visually responsive regions (PLV_v_) was also significant as was the partial correlation between both connectivity to visual regions and stP timescale and a neural populations ability to predict patient response time (RT) during the 1-back task.

The negative partial correlation between a neural population’s stP timescale and its functional connectivity to visually responsive populations throughout the brain was significant. This suggest that parts of VTC that communicate strongly with other visually responsive regions have shorter timescales. Furthermore, the negative partial correlations were significant between the magnitude of a neural population’s peak category-discriminant information and both its connectivity to visually responsive regions and those that were not. This shows that neural populations with stronger connectivity, especially to non-visual areas have less category-discriminant activity.

None of the measures of endogenous or stimulus-response timescale (prestimulus stP and stHFBB tau, initial rise duration, and maintenance) were significantly correlated with one another, with or without removing the effects of distance along the visual hierarchy (*Fig 6* and *Fig S5*). Thus, though there are gradients in neural timescales across VTC using each of these measures, neither these timescales nor their gradients are significantly correlated to one another even though they were measured from the same neural populations. This indicates that neural timescales are context dependent (prestimulus vs stimulus response, initial rise duration vs. maintenance, stP vs. stHFBB, etc. are all not significantly correlated) and measuring one type of timescale cannot be used to infer the general timescale of a neural population.

Partial correlations between nearly all the stimulus response variables (peak information, onset time, initial rise duration, and maintenance duration), other than the two timescales discussed in the previous paragraph (initial rise duration vs. maintenance duration), were significantly correlated with one another. This suggests that there are interactive factors driving these different aspects of the stimulus response.

The partial correlation between the ability of a neural population to predict RT and that neural population’s connectivity to visually responsive brain regions and the partial correlation between a neural population’s ability to predict RT and that neural population’s prestimulus stP timescale after removing the effect of distance along the visual hierarchy were both significant (*Fig. 6*). Thus, neural populations which integrate information over visual brain regions with short stP timescales were more predictive of behavior during the 1-back task observed here. This demonstrates that aspects of both local neural dynamics and long-range cortical interactions are intimately linked to a neural population’s role in visual perceptual behavior.

## Discussion

Taken together, these results illustrate interrelationships between a neural population’s anatomical location, its local dynamics, and its long-range functional connectivity, which ultimately influence that population’s role in perception. In the current study, progressing along the ventral visual hierarchy was associated with decreases in peak category-discriminant information, longer information onsets, longer durations of initial information processing, longer periods of information maintenance, longer prestimulus stP timescales but shorter prestimulus stHFBB timescales, and opposing changes in connectivity to visual and non-visual brain regions. These results suggest that the anatomical and physiological gradients that exist along the visual hierarchy influence almost all aspects of prestimulus and information processing dynamics, which may constrain how these neural populations process information and their computational role in cognition. Indeed, a subset of these functional gradients were correlated with the ability of a neural population’s activity to predict the speed of behavioral responses during a visual 1-back task. Furthermore, many aspects of stimulus response dynamics shared significant interrelationships with one another beyond any shared relationship with anatomical location. Functional connectivity was correlated to aspects of both the stimulus response and prestimulus timescales, demonstrating how long-distance interactions can influence local neurodynamics. However, prestimulus and poststimulus information processing timescales were not strongly correlated to one another, nor were the initial rise and maintenance of the visual response, suggesting that different aspects of neural dynamics arise through different processes and mechanisms.

Previous studies have observed that neural populations demonstrate longer timescales when moving from primary sensory and motor regions to association cortices (5, 8, 9, 12, 13, 18, 19). The increasing endogenous timescales of stP activity along the ventral visual hierarchy observed here further support this organizing principle of cortex. Notably though, the endogenous stHFBB timescales demonstrated the opposite relationship along VTC, with shorter timescales in more anterior parts of VTC. Furthermore, the timescale of the stP and stHFBB were uncorrelated, demonstrating a dissociation between the dynamics of these two signal components recorded from the same neural population. This highlights a need to better understand the differences in the physiological origins of stP and stHFBB signal components (29–31).

The duration that category-discriminant neural populations initially process category-selective information increased along the ventral visual hierarchy, which may be the result of increased computational demands involved in forming more complex and individuated representations in more anterior category-selective neural populations (32–35). However, in traditional models of perception, neural units are passive visual feature detectors, that either fire or not depending on the presence or absence of their preferred features (36). In these models, little difference should be seen in the speed that neural populations process information further downstream because these passive feature detectors, even if they are sensitive to complex features, should respond rapidly and automatically to the presence of that feature (36). In this study, the duration of the initial rise in visual responsiveness did not change along the hierarchy, which fits with these traditional models. However, the divergence in the duration of category-discriminant versus visual response dynamics does not fit with these models. Instead, these results support a model of ventral visual representations that evolve through time, with information processing dynamics governed by interactions between the information being processed locally and globally through long-range connections, which reflect top-down and recurrent interactions (27, 37–39).

Long-range functional connectivity demonstrated a crossover effect along the ventral visual hierarchy, with decreasing connectivity to visually responsive regions and increasing connectivity to those that were not, consistent with previous fMRI studies (1, 2). Some of these gradients in functional connectivity were also associated with gradients in neural timescales even after controlling for effects of distance along the visual hierarchy. Specifically, neural populations that were more strongly connected to visually responsive regions demonstrated shorter endogenous stP timescales. One potential explanation for this result is that neural populations which integrate primarily visual inputs have faster timescales compared to neural populations that have more diverse inputs so that they are prepared to rapidly process incoming visual information (2, 13, 25). Notably, the partial correlation between connectivity to regions that did not demonstrate strong visual responses and poststimulus stP timescale was not significant. Previous models have not investigated differential effects of long-range cortical interactions with visual versus non-visual regions on the timescale of neural populations (11). This may be an important consideration for future models. Given the variable coverage of brain regions across patients in the current study, future studies are necessary to tease apart the impact that connectivity with specific brain regions has on local cortical dynamics.

Neural populations that demonstrated higher peak category-discriminant neural activity had earlier onsets, shorter durations of initial rise, and maintained that information longer. Our simulations demonstrated that our measures of peak and duration are independent, confirming that this correlation is physiological and not an artifact of the analysis (*Fig. S6*). Longer initial rises in category-discriminant information with smaller peak information may reflect evidence accumulation over longer timescales in these neural populations (13). Whereas partial correlations between local neural dynamics and long-range cortical interactions demonstrates that, in addition to sharing strong gradients along the primary axis of sensory processing systems, these properties of neural populations are closely linked to each other. These links between local dynamics and long-range cortical interactions are likely conferred in part by shared biochemical, microstructural, and macrostructural connectivity gradients that exist along the ventral visual axis beginning early in cortical development (1, 2, 6, 40, 41).

Functional gradients in VTC were also correlated with the degree to which a neural population’s activity could predict perceptual behavior. In the current study, increased functional connectivity to visually responsive regions and shorter prestimulus stP timescales were associated with a greater ability for a neural population’s activity to predict RT in a 1-back task after removing the effect of distance along VTC. This suggests that these neural populations may play a larger role in the basic visual discrimination task studied here, though it is notable that these results cannot address relationships between neurodynamics and perception or behavior in other tasks. Behaviors involving more complex perceptual representations and/or more complex behavioral decisions may rely more heavily on neural populations with longer timescales and on higher order cortical regions (37, 38, 42–45). Future studies are required to determine if finer level of visual discrimination involving longer response times (46) reflect contributions from neural populations with different information processing timescales and functional connectivity patterns compared to those involved in the 1-back task studied here.

There were not significant correlations between stimulus response timescales and endogenous timescales, or between onset dynamics and maintenance dynamics. Different aspects of task-evoked timescales were not closely linked to one another, suggesting the physiological drivers of initial information processing and maintenance may be independent. Additionally, endogenous neurodynamic timescales did not generalize to stimulus related information processing timescales. Notably, this is unlike functional connectivity patterns, which were highly correlated across the stimulus response and prestimulus periods. The lack of significant correlation across the different measures of local neurodynamics highlights that endogenous neural timescales are not necessarily tightly related to task-evoked information processing dynamics (15, 47–49). Thus, inferences about a region’s computational role in cognition, including its temporal integration and segregation (10) or temporal response windows (18, 28), cannot be inferred from endogenous dynamics alone, as stimulus response and endogenous timescales are not necessarily strongly correlated. There is no single principle or process that governs a neural population’s timescales, e.g. timescales are not a static and inherent property of a neural population (10, 50). Rather, these results suggest that different kinds of timescales are governed by different combinations of factors that can depend on cognitive and neural context.

The current study highlights how large-scale anatomical and functional gradients interact to constrain local neural processing dynamics and computation. The anatomical gradients of dynamics and connectivity demonstrated here impose important constraints for future neurobiological models of visual perception. This architecture may help the brain achieve abstract and conceptual representations seen in more anterior VTC neural populations (32, 34, 35). While the present study examined these effects in visual processing, it is likely that similar principles apply to other hierarchically organized sensory and cognitive systems (2, 12, 14, 50).

Indeed, gradients in physiological, and thus functional, organization are likely in part conferred by corresponding gradients in growth factors and, in turn, gene expression during and persisting after cortical development (2, 7, 40, 41). Interactions among response properties and functional connectivity patterns of neural populations suggest that shared neurophysiological mechanisms tie large-scale and local processing dynamics together. Distinctions among and between endogenous and stimulus response timescales suggest that these neurodynamics are caused by distinct neurobiological mechanisms and play different roles in the brain. These results highlight the mutual interrelationships between a neural population’s position in the processing hierarchy, its functional connectivity, and its local dynamics, constraining its role in cognition.

## Acknowledgments

We would like to thank the patients and staff in the University of Pittsburgh Comprehensive Epilepsy Center at the University of Pittsburgh Medical Center for their participation in this research study. We would also like to thank Marlene Behrmann, Michael Tarr, and Max G’Sell for helpful suggestions in the design of the experimental analyses in addition to Michael Ward, Shawn Walls, and Ashley C. Whiteman for their help with data collection.

## Author contributions

Conceptualization, M.J.B. and A.S.G.; Methodology, M.J.B. and A.S.G.; Investigation, M.J.B., R.M.R., and A.S.G..; Formal Analysis, M.J.B.; Writing – Original Draft, M.J.B. and A.S.G.; Writing – Review & Editing, M.J.B., R.M.R., and A.S.G.; Resources, R.M.R. and A.S.G.; Funding Acquisition, A.S.G.;

## Data and materials availability

Data and analysis code will be made available upon request.

## Methods and Protocols

### Intracranial electroencephalography (iEEG) patients

Stereotactic depth and surface electrocorticography (ECoG) electrodes were implanted in ventral temporal cortex (VTC) of 41 patients (15 males, ages 19-65) for the localization of pharmacologically intractable epileptiform activity. Different aspects of these recordings from 38 of these patients were previously reported in (31). All patients gave written informed consent under protocols approved by the University of Pittsburgh’s Institutional Review Board. Electrode contacts that were identified as belonging to the seizure onset zone were not included in the analysis.

Electrodes were localized via postoperative CT scans or postoperative magnetic resonance images (MRI). Postoperative CT scans were co-registered to preoperative MRIs using Brainstorm (51). Surface electrode contacts were projected to the nearest reconstructed cortical voxel of the preoperative MRI scan to correct for brain-shift (52, 53). These electrode locations were then registered to the Montreal Neurological Institute (MNI) common space via patient-specific linear interpolations (54). VTC was defined as grey matter below the inferior temporal gyrus spanning from the posterior edge of the fusiform gyrus to the anterior temporal lobe in MNI common space.

Cortical distance between each electrode contact and the patient’s occipital pole was computed using the patient’s native neural anatomy. The occipital pole was defined as the intersection of the calcarine sulcus, inferior occipital gyrus, and superior occipital gyrus. The geodesic (cortical) distance between this point and the cortical surface coordinate nearest to each VTC electrode contact was computed using custom MATLAB scripts (55).

### Experimental Paradigm

All patients underwent a category localizer task containing images occupying approximately 6° x 6° of visual angle at the center of a stimulus display monitor positioned 2 meters from the patient’s eyes. Each stimulus was presented for 900 ms on a black background. Inter-stimulus intervals were 1500 ms with a random 0-400 ms jitter during which the patient saw a white fixation cross. Patients were instructed to press a button every time an image was presented twice in a row (1/6 of all trials). Repeat trials were excluded from further analysis. This left 70 trials per category to train and test the classifiers described in *Multivariate temporal pattern analysis*. Several patients underwent more than one run of this experiment and therefore had 140 or 210 trials per category. All experimental paradigms were presented via custom MATLAB scripts running the Psychophysics toolbox (56).

35 patients underwent a category localizer task consisting of pictures of bodies, faces, hammers, houses, words, and non-objects. Six patients underwent category localizer tasks with slightly different object categories but with identical stimulus parameters. One of these patients viewed pictures of bodies, faces, *shoes*, hammers, houses, and phase-scrambled objects. One viewed pictures of bodies, faces, *consonant-strings, pseudowords*, real words, houses and phase-scrambled objects. One patient viewed pictures of faces, bodies, *consonant-strings*, words, hammers, and phase-scrambled objects. One viewed pictures of faces, bodies, words, *pseudowords*, houses, and phase-scrambled objects. One viewed pictures of faces, bodies, words, *tools, animals*, houses, and phase-scrambled objects. One viewed pictures of faces, bodies, words, *tools, animals, numbers*, houses, and phase-scrambled objects.

### Intracranial recordings

Local field potentials were collected from iEEG electrodes via a GrapeVine Neural Interface (Ripple, LLC) sampling at 1 kHz. Notch filters at 60/120/180 Hz were applied online. Stimulus presentation was synchronized to the neural recordings via parallel port triggers sent from the stimulus displaying computer to the neural data acquisition computer. The signal was off-line filtered from 0.2-115 Hz using two-pass fourth order butter-worth filters via the FieldTrip toolbox (57). In addition to analyzing these single trial potentials (stP), we also extracted and analyzed the single trial high frequency broadband (stHFBB) activity of these electrodes, since these two components of the local field potential have been shown to contain complimentary information (24).

StHFBB activity was extracted via Morlet wavelet decompositions from 70-150 Hz over 200 ms Hanning windows with 10 ms spacing. The resulting power spectral densities were then averaged over these frequency components and normalized to a baseline period from 500 ms to 50 ms prior to stimulus onset to yield the stHFBB activity. Data was then epoched from -500 to 1500 ms around stimulus presentation. Trials during which the stP amplitude changed more than 25 microvolts across a 1 ms sample, or during which stPs exceeded an absolute value greater than 350 microvolts, or during which either the stHFBB or stPs deviated more than 3 standard deviations from the mean were all assumed to contain noise and were therefore excluded.

### Multivariate temporal pattern analysis (*Fig. 1, 2, & 3*)

Sliding, leave-one-out cross-validated, Gaussian Naïve Bayes classifiers were applied to 100 ms time windows with 10 ms stride to determine if stHFBB or stP recorded from individual VTC contacts contained category-discriminant information. The input to these classifiers was 100 ms (100 samples) of stP and 100 ms (10 samples) of stHFBB from a single electrode contact. The output of the classifier was the category of object presented during the corresponding trial. This procedure was repeated for all VTC contacts from time windows beginning at 100 ms prior to stimulus onset to 1000 ms after stimulus onset.

The category-discriminant information content within each neural population was estimated by computing the mutual information (*I(S’,S)*) between the output of the Gaussian Naïve Bayes classifiers (predicted category labels, *S’*) for a given 100 ms time window of neural activity and the actual presented stimulus (S):

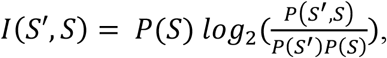

where *P(S’,S)* is the joint probability of the classifier correctly predicting the stimulus category *S* when the category was *S, P(S’)* is the proportion of times the classifier guessed a trial was of stimulus *S*, and *P(S)* is the proportion of trials which the stimulus presented was *S*. This allowed us to estimate the category-discriminant information contained within 100 ms time windows without estimating a joint probability table of neural responses that was intractable (58, 59). It has been shown that this estimate of information, which relies on a *P(S’,S)* derived by an external classifier and not the actual neural code, is an underestimate of the neural information content (60). Therefore, our calculated information is a lower bound for the actual neural information content.

Information content was averaged across all stimulus categories presented to the patient so as not to preclude electrode contacts as being selective for only one object category (61). A threshold for significant category-discriminant information was determined by randomly shuffling stimulus labels for a subset of VTC electrode contacts and repeating the same classification analysis 1,000 times for each electrode contact (62). Electrode contacts with the same number of runs of the category-localizer task demonstrated very similar null distributions and therefore we applied the result of this permutation test to all VTC electrode contacts. The threshold was chosen such that none of the random permutations for any electrode contacts in the subset reached the threshold, which corresponds to p < 0.001, corrected for multiple temporal comparisons. Electrode contacts with peak category-discriminant neural information exceeding this threshold were defined as category-discriminant.

We performed a similar decoding analysis to determine the time-course of visual responses in individual VTC contacts. This was done by classifying single trial baseline periods (100 ms to 0 ms prestimulus presentation) of neural activity from these neural populations against sliding 100 ms time-windows from -90 to 1000 ms post-stimulus presentation for all object categories treated as one class. This yielded a time-course of visual responses in each sampled neural population. By randomly permuting the label of the baseline versus evoked data and repeating the analysis in a subset of electrode contacts 1,000 times, we defined a threshold of visual information that no random permutation of the data achieved, corresponding to the p < 0.001, chance level, corrected for multiple temporal comparisons. We used this threshold to define visually responsive brain regions and those that were not, which were separated to calculate their differential contributions of functional connectivity to VTC electrode contacts with significant category-discriminant information.

### Estimating the dynamic properties of neural information processing (*Fig. 2* and *3*)

To estimate properties of the information processing dynamics of neural populations across VTC, the information time-courses derived from the Naïve-Bayes classifiers were first smoothed with a running average filter (width 50 ms). Next, onset latency of category-discriminant information was defined as the last time point that an electrode contact was below 10 % of the maximum information prior to the peak information. The initial rise in category-discriminant information was defined as the time between the onset and the point where the information time-course first exceeded 90 % of the peak information. These cutoffs were chosen to ensure that small deviations from chance-level information and peak information did not affect the estimated quantities. Our main findings were robust to specific choices in threshold (*Fig. S7*). Finally, we estimated the duration of information maintenance as the time between when the neural population first reached 90% of its peak information to when the neural population’s information first fell below 50% of this maximum after peaking. Similar dynamic properties were also estimated for visual response time-courses (*Fig. 3*) and information processing time-courses for specific object categories (*Fig. S3 & Table S1*).

### Defining category-selective VTC electrode contacts (*Fig. S3*)

To determine if neural populations with sensitivity to different object categories demonstrated differences in the gradients of their local dynamics or long-range functional connectivity, we isolated category-discriminant VTC neural populations that responded primarily to one object category. To do this we computed the event related potential and event related broadband responses to each category during the 1-back task. Next, any of the previously defined category-discriminant neural populations that contained maximum information to the same category that evoked the maximum response across either of these averages was classified as selective to that object category. We then characterized the information onset latency, slope, and connectivity of these neural populations using the procedures described above. For these analyses we used the category-specific information processing time-course derived from the Naïve Bayes classifiers prior to averaging over all categories in the main analysis.

### Information processing simulations (*Fig. S6*)

Simulations were used to test if increases in information processing duration exhibited along the ventral visual hierarchy could be explained by differences in peak information magnitude. Specifically, information time-courses were approximated as normal probability density functions (PDFs) parameterized by a mean, standard deviation, and magnitude (constant scaling). Normally distributed noise with the same standard deviation as prestimulus (−400 to 0 ms) information in category-selective VTC electrode contacts was then added to these curves. 1000 simulated signals were computed for each different PDF magnitude and standard deviation.

Information processing duration was calculated using the same procedure described for the actual signal, by calculating the time between when the signal first reached 90 % of its maximum amplitude and the last time it was below 10 % of its maximum before that. We then calculated the Spearman correlation between information processing duration when varying the PDF’s standard deviation (to mimic changes in slope of the information processing time-course) and when varying the information’s peak amplitude. Peak amplitude was varied from the minimum to maximum peak information in category-selective VTC electrode contacts. During the simulation investigating the effect of slope on information processing duration, signal amplitude was fixed at the average peak information in category-selective VTC electrode contacts.

### Characterizing endogenous neural timescales (*Fig. 4*)

The endogenous timescales of VTC populations were characterized by computing the autocorrelation of prestimulus (−500 ms to stimulus onset) stPs and stHFBB activity from 1-250 ms lags during each clean trial of the 1-back task. These prestimulus autocorrelation functions were then averaged over all trials. The average autocorrelation function for each electrode contact was then fit with a single exponential decay function:

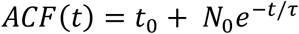

The neural timescale (tau), which measures the rate at which the autocorrelation function decays, was then correlated with several other functional properties of the neural population. This estimation of neural autocorrelations and computation of tau is similar to the procedure described in (5).

### Functional Connectivity (*Fig. 5*)

To determine the connectedness of VTC neural populations to the rest of the brain, phase-locking values (PLVs) were calculated between neural populations with above chance levels of category-discriminant information and all other electrodes within the same patient (regardless of category-discriminant information content). Electrode contacts within 1 cm of the category-discriminant electrode were not included in the analysis to rule out effects caused purely by volume conduction. PLVs measure instantaneous phase-coupling across different brain regions independent of differences in amplitude, unlike coherence metrics (63). This makes PLVs more sensitive to detecting weakly coupled oscillators despite differences in amplitude (64). This coupling of oscillations is thought to indicate event-related communication between electrode contacts.

The instantaneous phase of each electrode contact during all category-localizer trials was computed via convolution of the filtered neural activity (from 1-115 Hz) with Morlet wavelets of frequencies ranging from 1-60 Hz (width = 5). This convolution allowed the separation of signal phase from envelope at each frequency (65). Next, the PLV was computed by taking the vector average of the phase difference between two electrode contacts at each time point. PLVs close to 1 indicate two electrode contacts have similar phase differences at this frequency and time point across all trials. Conversely, if this number is close to 0, the phase difference between these electrode contacts is random at this given frequency and time point.

A spectral window of interest was defined to capture the part of the PLV spectrogram that showed increased functional connectivity across all category-discriminant VTC neural populations. We chose to focus on the time windows from -450 to 0 ms before stimulus onset to capture prestimulus functional connectivity of the neural populations. Next, we determined which frequency components demonstrated increased stimulus-evoked functional connectivity across VTC. To do this we averaged the PLVs from 50 to 500 ms and performed a paired t-test against the average PLV from -450 ms to 0 ms before stimulus presentation between the category-selective VTC electrode contacts and the rest of the electrode contacts in the same patient. This analysis revealed that frequency components between 1 and 22 Hz all had significantly greater phase-locking across all category-discriminant VTC electrode contacts relative to baseline on average from 50 to 500 ms after stimulus presentation (p < 0.001, corrected).

Therefore, we averaged the PLVs across electrode contacts from 2 to 22 Hz (discarding 1 Hz frequency band to increase the temporal precision of our estimated phase-locking), and -450 to 0 ms before stimulus onset to calculate the functional connectedness of these same regions. We separately averaged the connectivity of category-discriminant VTC neural populations with visually responsive regions (defined above) and those that were not to determine if there were connectivity differences across these neural populations. Average functional connectivity from - 450 to 0 ms prestimulus and 50-500 ms after stimulus presentation were strongly correlated with one another (ρ = 0.96, p < 0.001). Thus, results do not substantially change if either the prestimulus or post stimulus PLV is used.

### Predicting patient response time from category-selective VTC (*Fig. 6*)

To test for differences in the correlation between category-selective VTC population activity and behavior, patient RT was predicted using the neural activity from each category-selective contact. Specifically, a sliding window L2-regularized multiple regression (100 ms window, 10 ms stride) was used to predict patient RT from stP and stHFBB activity using a leave-one-trial-out cross-validation procedure. Only trials when the patient correctly reported that an object was repeated twice in a row were included in the analysis. The maximum Spearman correlation between the patient’s RTs and the sliding-window RT predictions from 0-1000 ms after stimulus presentation was considered as the neural population’s correlation with behavior. This correlation was then correlated with that population’s dynamics, connectivity, and anatomical location.

### Statistics

Spearman rank-order correlations were used to calculate the correlations between anatomical position and aspects of the neural information time-courses calculated above. Spearman rank-order partial correlations were used to calculate the correlation between variables while correcting for correlations shared with other variables. Benjamini-Hochberg False Discovery Rate estimation which is valid for dependent hypothesis tests was used where noted (66). Paired T-tests were used to determine if there were differences in the dynamics of processing different levels of information (visual vs. category-discriminant) in the same electrode contacts.

Rank-order mixed-effects models were used to control for random effects of cross-patient variability while examining the main effects of connectivity and anatomical position on information processing dynamics. We chose to fit these mixed-effects models with equal slopes but random intercepts across patients to ensure the models converged. Because observations in mixed-effects models are not independent, it is difficult to determine the appropriate degrees of freedom. This makes estimation of p-values impossible without appropriate approximation. Therefore, to derive p-values for the main effects of the mixed-effects models, we use the Satterthwaite approximation, which has been shown to produce acceptable Type 1 error rates with relatively few samples (67).

Linear multiple regression models were used to compare gradients of information processing in VTC neural populations that were selective for different object categories. We only included the categories that most patients saw (bodies, words, faces, hammers, houses, and phase-scrambled objects). Specifically, linear models were used to predict information onset latency, peak, processing duration, maintenance duration, and connectedness as a function of the category-selective neural populations’ distance from the occipital pole with an added factor indicating which category the neural population was selective for (*Fig. S3*). Linear mixed-effects models were initially used for this analysis to simultaneously control for random effects across patients. However, these models failed to converge, likely indicating an insufficient number of data points per category and patient to estimate these random effects. Because face-selective electrode contacts were most prevalent in our population we used this as our baseline and compared all other categories to face-selective electrode contacts (*Table S1*). Analysis of covariance was also used to determine if there was a significant difference in information processing gradients or connectedness across hemispheres (*Supplementary Text*).

