## Supplementary Materials for "Interacting cortical gradients of neural timescales and functional connectivity and their relationship to perceptual behavior"

**Extended Data:**

Supplementary Text

Figures S1-S7

Table S1

### Supplementary Text

Interhemispheric differences in ventral stream information processing dynamics.

Given asymmetries in anatomical organization and response tuning across hemispheres (68), we sought to determine if there were differences in information processing or functional connectivity gradients in left versus right VTC. We found 263 electrode contacts with significant category-discriminant information in left VTC and 127 in the right. The disparity in left and right hemisphere contacts is similar to the different proportion of total contacts implanted in the left versus right VTC across patients (1,258 in left, 698 in right). Using these category-discriminant electrode contacts, we ran an analysis of covariance to determine if there was an interaction between anatomical position and hemisphere when predicting information processing dynamics or functional connectedness. There was a significant interaction between cortical distance along the visual hierarchy and hemisphere when predicting onset latency, indicating a more dramatic gradient in the right compared to left hemisphere ($F(1) = 16.19$, $p < 0.001$), and connectivity to regions that were not visually responsive, indicating a more dramatic gradient in the left compared to right hemisphere ($F(1) = 11.76$, $p < 0.001$). There were no significant interactions between hemisphere and distance along the visual hierarchy in predicting peak information ($F(1) = 1.25$, $p = 0.26$), average PLV to visually responsive regions ($F(1) = 0.44$, $p = 0.51$), or duration of initial rise in information ($F(1) = 0.91$, $p = 0.34$). The interaction between hemisphere and distance along the visual hierarchy was trending when predicting information maintenance duration ($F(1) = 4.49$, $p = 0.035$, uncorrected). This suggests that there were differences in anatomical gradients of information onset and connectivity with regions that were not visually responsive across hemispheres in our sampled neural populations.

Differences in functional anatomical gradients for different object categories

It has previously been suggested that VTC circuits responsible for processing different categories of objects exhibit different processing dynamics related to the rate at which those objects are encountered during natural vision (69). To investigate differences in information dynamics between neural populations tuned primarily to single object categories, we identified category-discriminant VTC electrode contacts that demonstrated maximum evoked responses to the object category which also had the most information (see *Methods*). This procedure revealed that 246 of the 390 category-discriminant VTC neural populations were predominantly selective for a single object category: 66 were selective for faces, 50 for words, 47 for houses, 31 for phase-scrambled objects, 21 for bodies, 20 for hammers, 7 for tools, 3 for pseudowords, and 1 for consonant-strings (*Fig. S3*). Next, we fit linear multiple regression models to explain information processing dynamics of the neural populations that were selective to the object categories that most patients saw (bodies, faces, words, hammers, houses, and phase-scrambled objects) as a function of the neural population's distance from the occipital pole and the category it was selective for (see *Methods*). This allowed us to compare the functional anatomical gradients specific to different categories of objects in VTC.

The main effects revealed by this procedure are contained in *Table S1* and *Fig. S3*. The information onset of face-selective neural populations increased at a rate of 11 ms per cm traveled along VTC. Neural populations selective for all other categories demonstrated faster increases in onset latency along this axis compared to face-selective neural populations. Peak face-selective information was not significantly different when moving along the ventral visual hierarchy. However, peak information for the other observed categories decreased faster along this axis. There was no significant change in the duration of the initial rise of face-selective information when moving up the visual hierarchy, nor was the gradient for any category significantly different from faces. There was also no significant change in the duration that face-selective information was maintained in neural populations further up the visual hierarchy, but information was maintained for shorter durations moving along this axis for all other object categories.

Functional connectivity to visually responsive regions did not significantly decrease along the ventral visual hierarchy when looking at face-selective information. However, visual connectivity decreased more quickly when moving along VTC in word-, hammer-, and house-selective populations compared to face-selective populations. Finally, there was not a significant gradient in the functional connectivity of face-selective populations to regions that were not visually responsive along the visual hierarchy, nor were there any significant differences observed for the other object categories.

In summary, at the level of individual categories, face-selective neural populations demonstrated faster onsets of information processing, larger peaks, longer durations, and increased connectivity to visually responsive neural populations compared to the other object categories. These differences in information processing dynamics across category-selective neural populations is consistent with previous fMRI studies demonstrating different preferential rates of stimulus presentation for different categories of objects (69). Together, these results suggest differences in information processing dynamics for different categories of objects, which may be related to the functional interactions that facilitate their recognition or how these stimuli are naturally encountered in the real world (69).

Comparing neuroanatomical gradients exhibited in single trial potentials versus high frequency broadband activity

Previous studies have identified differences in the information contained within single trial potentials (stP) and single trial high frequency broadband activity (stHFBB) (29, 31) and others have suggested that these components of have different physiological generators (30, 70). To investigate the degree to which neuroanatomical gradients in neural dynamics and long-range functional interactions were consistent across these signal components, we re-ran the main analyses of this study using stP and stHFBB activity separately. We isolated 380 electrode contacts that demonstrated above-chance category-discriminant activity ($p<0.001$, permutation test) in their stP and 150 contacts that were discriminant in their stHFBB activity (*Fig. S4*).

In contacts selective in their stP, gradients in the onset, duration of initial rise, peak magnitude, and maintenance duration of category-selective information were consistent with those identified when jointly classifying stP and stHFBB activity. The gradients in stP timescale as well as connectivity to visually responsive regions and connectivity to regions that were not visually responsive were also consistent. Unlike the jointly classified data, electrodes selective in their stP demonstrated a decreasing ability to predict patient RT along the visual hierarchy ($\rho(380) = -0.11$, $p = 0.026$, uncorrected).

In contacts selective in their stHFBB activity, the correlations between local neural dynamics and long-range functional connectivity were smaller but were mostly consistent with those observed on the jointly classified data. However, connectivity to regions that were not visually responsive had the opposite relationship with distance along VTC compared to the jointly classified contacts ($\rho(150) = 0.30$, $p < 0.001$). Additionally, contacts that were sensitive to object category in their stHFBB activity demonstrated increasing ability to classify patient RT from the stHFBB activity when moving up the visual hierarchy ($\rho(150) = 0.24$, $p = 0.0026$). These results illustrate that functional anatomical gradients in information processing dynamics in ventral visual cortex are largely consistent across stP and stHFBB activity; however, these signal components do demonstrate differences in their anatomical gradients in timescales, connectivity to regions that are not visually response, and ability to predict patient response time during a visual 1-back task.

**Fig. S1.**

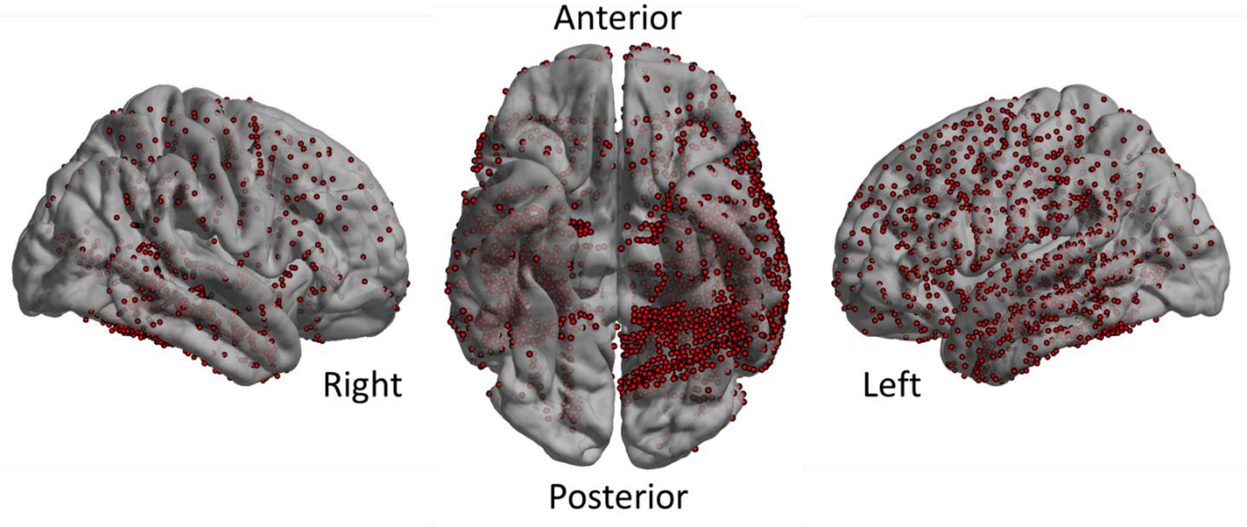


Intracranial electrode contact coverage of 35 patients with category-discriminant VTC electrode contacts. Contacts that appeared to be outside of the MNI standard brain due to differences in individual brain sizes were projected to the nearest MNI cortical vertex in this figure solely for illustrative purposes.

**Fig. S2.**

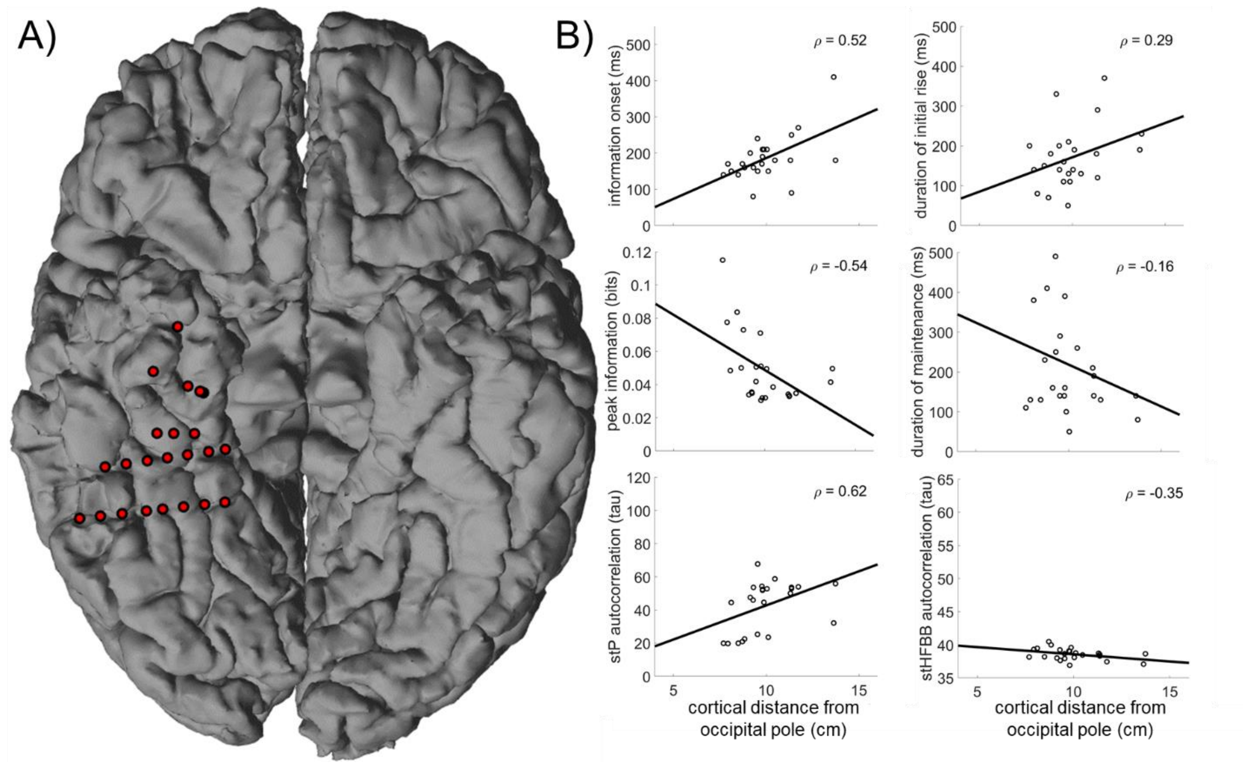


Information processing and neural timescales in VTC of one patient. A) Spatial topography of VTC electrode contacts with above chance ($p < 0.001$, corrected for multiple temporal comparisons) category-discriminant information on the individual's anatomy. B) Relationship between information processing dynamics and neural timescales examined in *Fig. 2* and *3* and the cortical distance along VTC. Spearman correlation coefficient displayed in the top right of each panel. Line represents least-squares approximate fit. Information onset, peak information, and stP timescale (tau) were all significantly correlated with distance along VTC with the same direction as the group-level gradients. Other properties of the neural population's dynamics demonstrated similar gradients with distance along VTC as the group-level data but were not significant across these 24 electrodes.

**Fig. S3.**

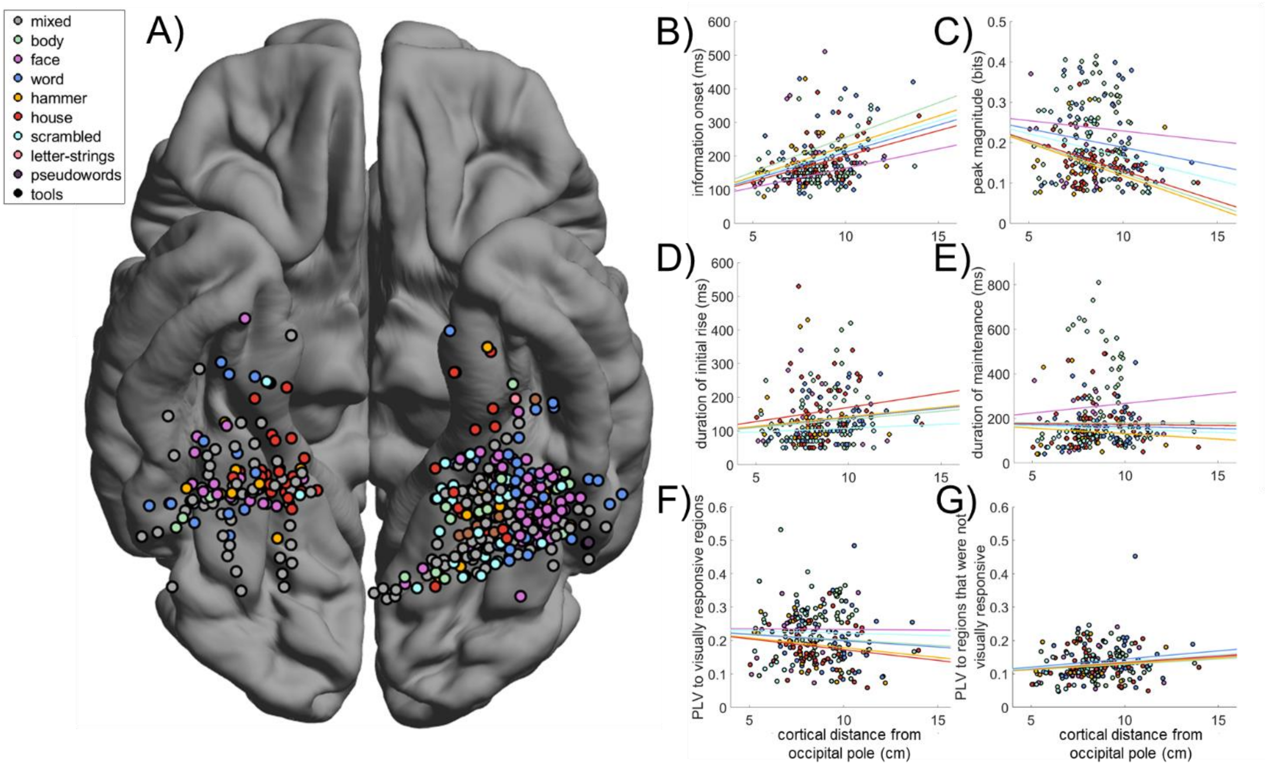


Differences in functional anatomical gradients across neural populations selective for different object categories. A) Spatial topography of electrode contacts predominantly selective for one object category. Grey electrode contacts contained a significant amount of category-discriminant neural information but were sensitive to more than one image category. B) Onset of category-discriminant information processing as a function of cortical distance along the ventral visual hierarchy for neural populations selective for faces, bodies, words, hammers, houses, and phase-scrambled objects. Lines are derived from multiple linear regression analyses. Corresponding statistics are contained in *Table S1*. Face-selective information demonstrated faster propagation along the visual hierarchy compared to the other object categories. C) Relationship between the duration of the initial rise in category-selective information versus distance along VTC. There was no significant difference in this gradient between neural populations selective for different object categories. D) Peak category-discriminant information as a function of distance along the visual hierarchy for VTC neural populations selective for different object categories. Face-selective information decayed less when moving down the ventral visual hierarchy compared to the other object categories. E) Maintenance duration of category-selective information as a function of distance along VTC. Face-selective neural populations demonstrated the greatest increase in information maintenance duration when moving along VTC compared to the other object categories. F) Relationship between connectedness to visually responsive regions and distance along the visual hierarchy for VTC neural populations selective for different object categories. Word-, hammer-, and house-selective neural populations demonstrated greater decreases in connectivity to visually responsive regions when moving along VTC compared to face-selective neural populations. E) Relationship between connectedness to regions that were not visually responsive and position in the visual hierarchy for VTC neural populations selective for different object categories. There was no significant difference in this anatomical gradient across neural populations selective for different object categories.

**Table. S1.**

| | Estimate | St. Error | T-stat | Estimate | St. Error | T-stat |
|---|---|---|---|---|---|---|
| | **Information onset (ms/cm)** | | | **Duration of initial rise (ms/cm)** | | |
| **Faces** | **11.46** | **2.83** | **4.04*****| 5.42 | 3.38 | 1.60 |
| **Bodies (difference from faces)** | **9.15** | **1.98** | **4.61***** | -0.56 | 2.37 | -0.24 |
| **Words (difference from faces)** | **4.72** | **1.37** | **3.46***** | 0.01 | 1.63 | 0.01 |
| **Hammers (difference from faces)** | **6.52** | **2.06** | **3.16**** | 0.22 | 2.46 | 0.09 |
| **Houses (difference from faces)** | **3.60** | **1.43** | **2.51*** | 2.99 | 1.71 | 1.75 |
| **Scrambled (difference from faces)** | **5.56** | **1.72** | **3.23**** | -3.18 | 2.06 | -1.55 |
| | **Peak information magnitude (bits/cm)** | | | **Duration of maintenance (ms/cm)** | | |
| **Faces** | -0.0051 | 0.0035 | -1.48 | 8.64 | 5.76 | 1.50 |
| **Bodies (difference from faces)** | **-0.0105** | **0.0024** | **-4.32***** | **-8.73** | **4.03** | **-2.17*** |
| **Words (difference from faces)** | **-0.0041** | **0.0017** | **-2.419*** | **-10.39** | **2.77** | **-3.74***** |
| **Hammers (difference from faces)** | **-0.0111** | **0.0025** | **-4.39***** | **-13.53** | **4.19** | **-3.23**** |
| **Houses (difference from faces)** | **-0.0098** | **0.0018** | **-5.59***** | **-9.44** | **2.91** | **-3.24**** |
| **Scrambled (difference from faces)** | **-0.0064** | **0.0021** | **-3.04**** | **-11.57** | **3.50** | **-3.3017**** |
| | **Connectivity to significantly visually responsive regions (a.u.)** | | | **Connectivity to regions that were not significantly visually responsive (a.u.)** | | |
| **Faces** | -0.0004 | 0.0033 | -0.12 | 0.0036 | 0.0020 | 1.79 |
| **Bodies (difference from faces)** | -0.0031 | 0.0023 | -1.32 | -0.0004 | 0.0014 | -0.28 |
| **Words (difference from faces)** | **-0.0034** | **0.0016** | **-2.09*** | 0.0012 | 0.0010 | 1.24 |
| **Hammers (difference from faces)** | **-0.0055** | **0.0024** | **-2.24*** | -0.0001 | 0.0015 | -0.10 |
| **Houses (difference from faces)** | **-0.0060** | **0.0017** | **-3.57***** | 0.0002 | 0.0010 | 0.20 |
| **Scrambled (difference from faces)** | -0.0011 | 0.0020 | -0.54 | 0.0005 | 0.0012 | 0.38 |

Statistically evaluating differences in functional anatomical gradients across processing hierarchies for different categories of objects. This table summarizes the statistical effects of the models illustrated in *Fig. S3B-G*. Contacts with mixed selectivity were not included in the models. Face-selective neural populations were used as the base-level of the analyses because they were the most prevalent. Therefore, all other rows indicate the difference between the slope of the gradient in face-selective versus other category-selective populations. There were significant differences in the anatomical gradients of information onset, peak, duration of maintenance, and connectivity to visual neural populations across neural populations selective for different object categories. Significant effects are highlighted in bold. (* $p < 0.05$, ** $p < 0.01$, *** $p < 0.001$)

**Fig. S4.**

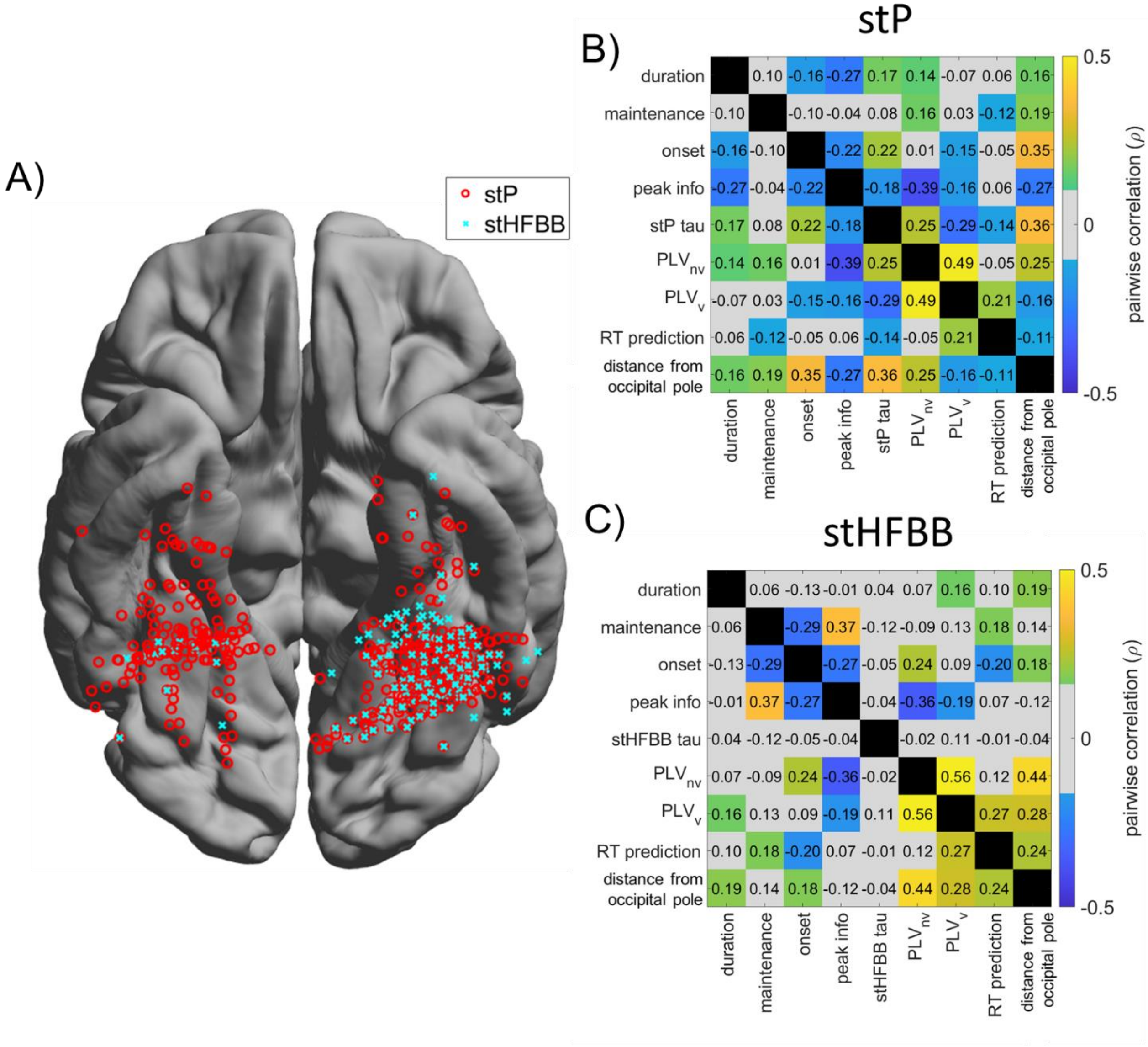


A) Electrode contacts demonstrating above-chance levels of category-discriminant information in single trial potentials (stP; n = 380) and single trial high frequency broadband (stHFBB; n = 150) when these signal components are classified separately. B) Pairwise Spearman correlations (ρ) between gradients in the corresponding row and column computed using electrode contacts selective in stP. Information processing metrics (onset, peak, maintenance, and rise durations) were computed for decoding time-courses derived from each signal separately. Shading indicates strength and direction of pairwise correlation. Grey squares indicate pairwise correlations that were not significant at the $p < 0.05$ level, uncorrected. The false-discovery rate adjusted critical value was estimated to be $\rho = \pm 0.138$. C) Same as panel B, but for contacts selective in their stHFBB activity. The false-discovery rate adjusted critical value was estimated to be $\rho = \pm 0.269$. The pairwise correlation for jointly classified stP and stHFBB data is available in *Fig. S5*.

**Fig. S5.**

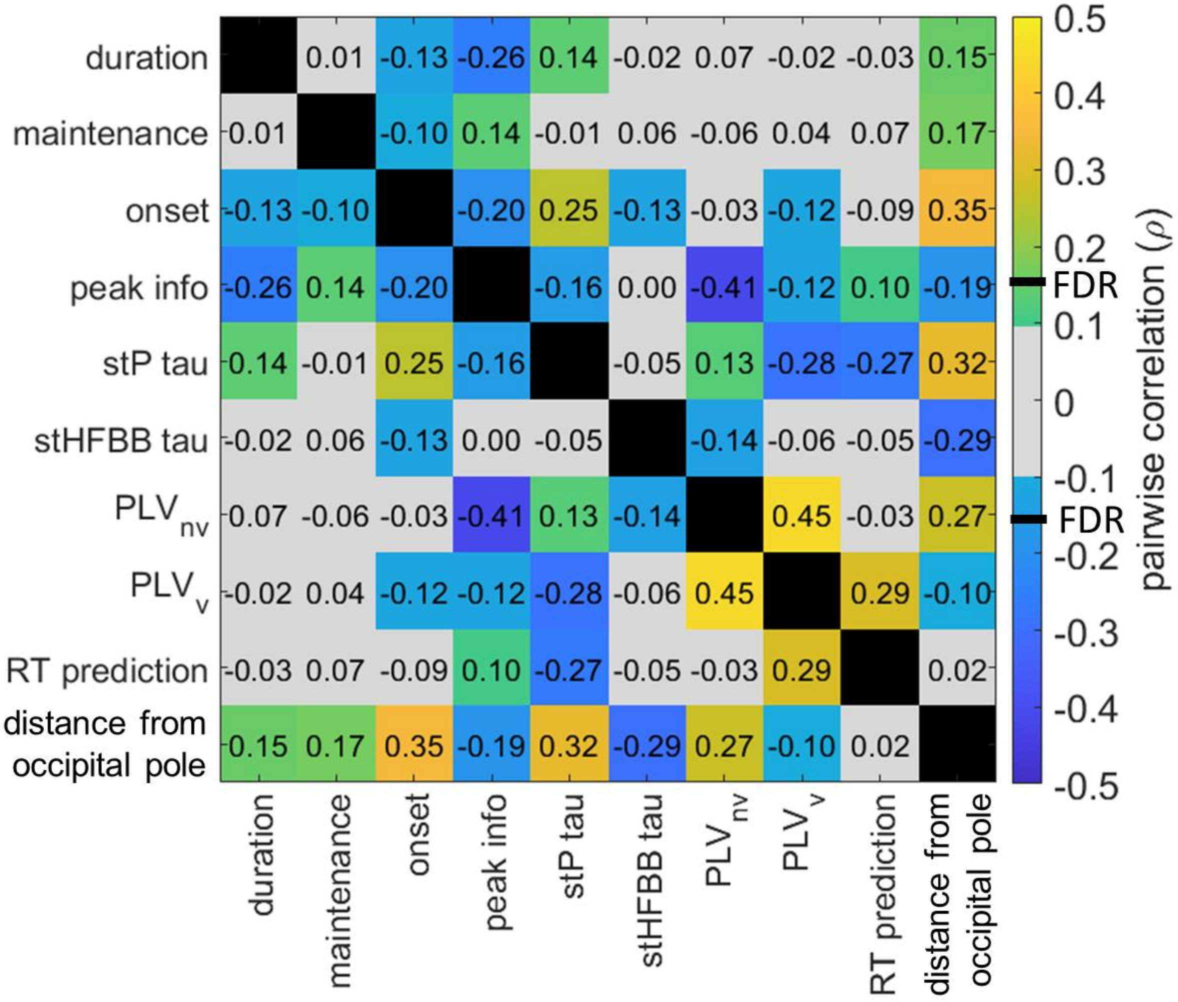


Pairwise correlations between dynamic and functional properties of VTC neural populations (full correlation matrix, no partial correlations). Within each box is the pairwise correlation (ρ) between the variables in the corresponding row and column, like *Fig. 4*, without removing the shared correlations with distance along VTC. Shading indicates strength and direction of pairwise correlation. Grey squares indicate pairwise correlations that were not significant at the $p < 0.05$ level, uncorrected. The false-discovery rate adjusted critical value was estimated to be $\rho = \pm 0.154$.

**Fig. S6.**

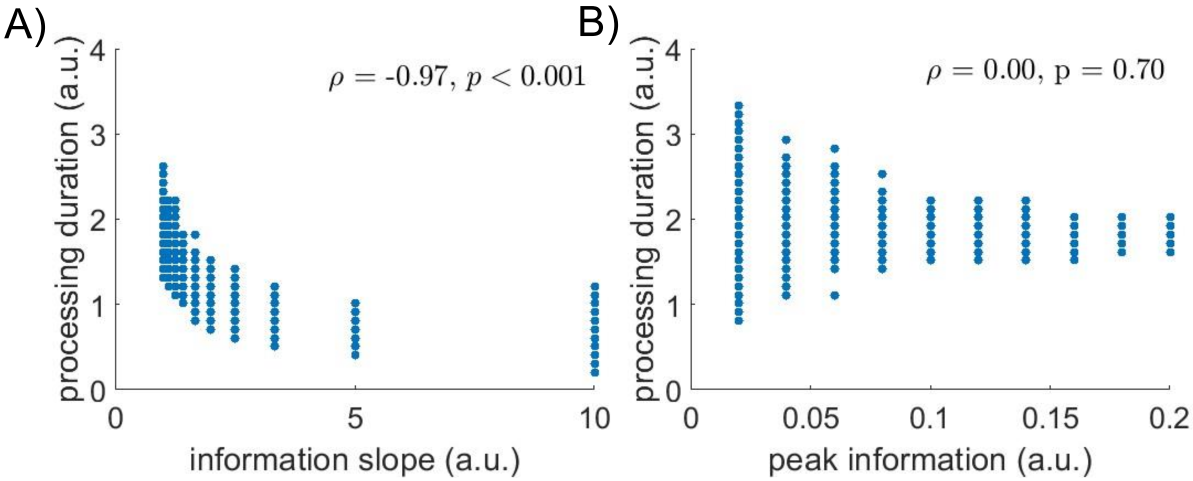


Simulating the effects of changes in information slope and peak amplitude on information processing duration. Information time-courses were simulated using a normal probability density function with similar signal to noise properties as category-selective information time-courses obtained from VTC. A) The slope of the information time-course was strongly correlated with the duration of information processing as expected. B) The duration of information processing was not correlated with changes in peak information. These simulations support that increases in information processing duration along the ventral visual hierarchy is not driven by differences in peak information amplitude.

**Fig. S7.**

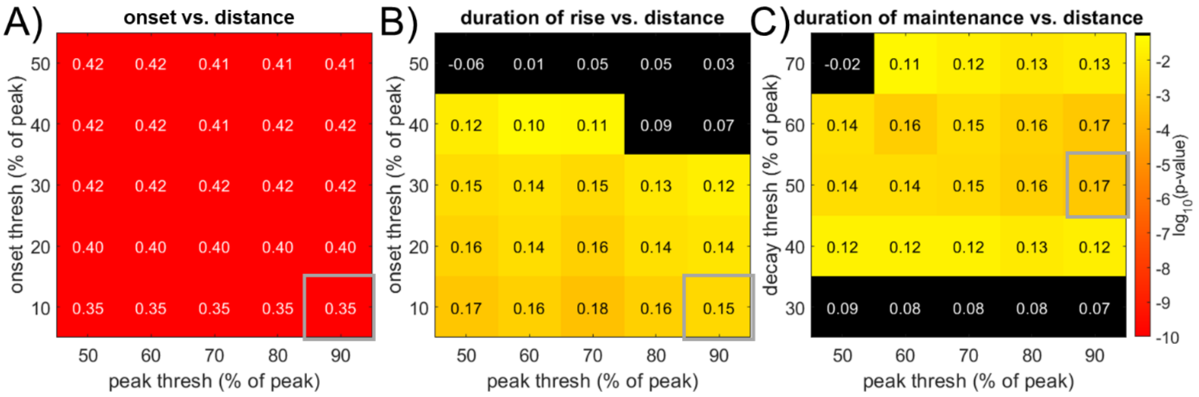


Anatomical gradients in information processing dynamics are robust to choices of onset and peak threshold. A) Spearman's correlation between a neural population's cortical distance along the visual hierarchy and its onset latency derived using different cutoff thresholds for onset and peak. Color of each square indicates p-value (color-bar on the right, log-scale). Inset of each square is the corresponding Spearman's ρ (n = 390). Gray square indicates result reported in the main text. Correlations derived from all criteria are very strong. B) Correlation between the duration of the initial rise in category-discriminant information and distance along the visual hierarchy with different onset and peak thresholds. Black squares indicate p-values greater than 0.05 uncorrected. C) Correlation between the duration of information maintenance and distance along VTC using different peak and decay thresholds. Decay was measured as the time between the reaching the peak threshold and the time the signal decayed below the decay threshold after reaching its absolute peak.
